# A chaotic viewpoint-based approach to solve haplotype assembly using hypergraph model

**DOI:** 10.1101/2020.09.29.318907

**Authors:** Mohammad Hossein Olyaee, Alireza Khanteymoori, Khosrow Khalifeh

## Abstract

Decreasing the cost of high-throughput DNA sequencing technologies, provides a huge amount of data that enables researchers to determine haplotypes for diploid and polyploid organisms. Although various methods have been developed to reconstruct haplotypes in diploid form, their accuracy is still a challenging task. Also, most of the current methods cannot be applied to polyploid form. In this paper, an iterative method is proposed, which employs hypergraph to reconstruct haplotype. The proposed method by utilizing chaotic viewpoint can enhance the obtained haplotypes. For this purpose, a haplotype set was randomly generated as an initial estimate, and its consistency with the input fragments was described by constructing a weighted hypergraph. Partitioning the hypergraph specifies those positions in the haplotype set that need to be corrected. This procedure is repeated until no further improvement could be achieved. Each element of the finalized haplotype set is mapped to a line by chaos game representation, and a coordinate series is defined based on the position of mapped points. Then, some positions with low qualities can be assessed by applying a local projection. Experimental results on both simulated and real datasets demonstrate that this method outperforms most other approaches, and is promising to perform the haplotype assembly.

## Introduction

Improving the high-throughput DNA sequencing technologies dramatically decreased the costs of genome sequencing methods. This achievement help researchers to understand the variation of individual’s genomic data and pave the way toward individualized strategies for diagnostic or therapeutic decision-making [1]. The most frequent type of genetic variation is the single nucleotide polymorphisms (SNPs). Each SNP is just a mutation over similar distinctive positions on the DNA sequences of homologous pair of chromosomes in an individual, and among the corresponding DNA sequences of the whole population. Similarly, the term “allele” refers to different forms of a gene at one loci. Accordingly, four different alleles are possible for a given SNP site. Nonetheless, most SNPs are bi-allelic containing only two kinds of alleles, which can be simply denoted by ‘0’ and ‘1’[2]. Each SNP contains valuable information about genomic alternations. Experimental studies revealed that SNPs have been clustered across the human genome and are not randomly distributed [3]. In line with this assumption, linkage disequilibrium (LD), demonstrates that there are correlations and spatial dependencies among neighboring SNPs. Different SNPs on the string of DNA is known as a haplotype. In other words, a haplotype could be considered as the combinations of marker alleles which are positioned closely together on the same strand of DNA, and tend to be inherited together from parents to offspring [4]. It has been shown that some diseases such as sickle-cell anemia [5], cystic fibrosis [6] and hemochromatosis [7] are more common in specific ethnic populations due to unique genetic mutations in their genomes; but they are rarely found in others. There are also reports indicating that different populations may have various responses to drugs. [8–10]. These findings demonstrate that haplotypes in human genomics data could be a useful and informative tool in mapping genes that are involves in representative diseases, as well as personalized medicine [11]. Haplotypes can also be used to investigate the pattern of inheritance over evolution, human migration, and the genetically aspects of populations [12–14]. Genetic association analysis for gene mapping can also be improved by haplotype analysis [15]. Also, it is possible to detect errors and missing sequencing data in experimental sequencing of DNA sequences using the information of haplotypes [16].

It is worth mentioning that the experimental analysis of haplotypes is labor-intensive and expensive. Moreover, it can be used only for constructing local haplotypes. In other words, human haplotypes are provided as sequencing reads or fragments. It is a vital task to obtain haplotype information from the numerous fragments due to its profound impacts on different aspects of medicine and molecular biology[15,17–19]. However, the detection of genetic variations has critical limitations compared with the molecular approaches. According to the type of input data, the existing methods of haplotype reconstruction are divided into two main categories, including single individual haplotyping (SIH) and haplotype inference. SIH methods receive several fragments that have been sequenced from a given chromosome. It is to be noted that most of the fragments contain gaps, and are usually disrupted by noise. To cope with these problems, the input fragments are clustered based on their similarities. Then, the haplotypes can be reconstructed using the center of each cluster [4]. The haplotype inference methods receive genotype information of several individuals as input data and infer their related haplotype sequences [20]. It is worth noting that each genotype represents a combination of haplotypes on the homologous chromosomes.

With increasing the size of data, a growing number of researchers have tried to solve haplotype assembly problem. Moreover, several computational models, including minimum fragment removal (MFR), minimum error correction (MEC), minimum SNP removal (MSR), and the longest haplotype reconstruction (LHR), have been developed to cope with the SIH problem. The MEC is one of the most popular and successful algorithms compared with the models as mentioned above [4,21–28]. This model attempts to cluster the input fragments, such that all the fragments belonging to a specified cluster to be compatible. Otherwise, they will be compatible by applying the minimum alternations. The current approaches can be divided into exact and heuristic methods. Since finding the optimal minimum error correction is NP-Hard, the exact approaches have exponential complexity [21]. Among exact solutions, WhatsHap[29] is regarded as a pioneering method, which is dynamic programming-based and utilizes a weighted variant of the MEC. The experimental results demonstrate that it can process long reads at coverage up to 20×. In[30], the authors proposed a parallel version of WhatsHap which is able to process higher coverages up to 25×. AROHap [24] is a recently published evolutionary-based method that exploits the asexual reproduction optimization algorithm to solve the SIH problem. In this method, the fitness function is designed based on the MEC model. In [26], a heuristic method, namely, Fasthap was developed, where it makes a weighted fuzzy conflict graph based on the MEC model. Furthermore, the constructed graph is used to cluster the input fragments. Fuzzy C-means (FCM) approach has been applied in [25] to enhance the performance of the proposed method in clustering the fragments. However, this method obtains low performance in dealing with noisy fragments. Some popular methods, including MCMC [31], HapCUT [27], and HapCUT2 [32], have differently construct the graph. These methods start with a set of arbitrary sequences as initial haplotypes, and improve it step by step concerning the input fragments. They make a similar weighted graph in their distinctive model. However, instead of fragments, SNPs are used as vertices of the graph. Each pair of SNPs is connected if they are covered by at least one input fragment. The weight of each edge determines the amount of consistency with their corresponding positions in the current haplotypes. Although this model efficiently determines the consistency of the current haplotype with the input fragments, the existing gaps and noise lead to a loss of accuracy in determining the weight of edges. In [33]. It has been proved that the hypergraph can precisely describe the distance of input fragments.

Although, various methods have been developed to solve the SIH problem, most of them can only be applied to diploid organisms, and fail to consider polyploid organisms. It should be noted that the haplotype reconstruction in polyploid type is more complicated than a diploid one. Suppose that *P* is the number of ploids, and *m* is the length of haplotype sequences. In this case, there are at least 2^*m*−1^(*P* − 1)^*m*^ different solutions for phasing the haplotypes [23]. Recently, several studies, such as [34], [23], [35], and [36], have been conducted on the polyploid organism. Althap[23] and SCGD[36] are two recently developed methods based on matrix factorization to solve the SIH problem. H-PoP [34] is a heuristic method that divides the input fragments into P clusters. Therefore, the members of each cluster have the minimum distance with each other and are entirely far from the fragments of other clusters. Belief propagation (BP) [35] is another method addressing the SIH problem by mapping the MEC model to a decoding mechanism. It involves a message transmission in a noisy channel. In this context, it has been reported that the haplotype’s blocks with proper lengths can exhibit chaotic behavior. This feature has been recently used to improve the reconstruction rate in the single individual haplotyping problem[37].

Considering the chaotic nature of haplotype sequences, in this paper, an iterative algorithm is proposed to reconstruct the haplotypes using the hypergraph model. The method includes two main steps. Firstly, an iterative mechanism is applied due to the SNP matrix to construct the haplotype set, and the consistency between SNPs is modeled based on the hypergraph. Then, the corrected parts of the haplotypes are determined by partitioning the hypergraph.

This step is followed by transforming the obtained haplotypes into a line using the chaos game representation, where a coordinate series is defined based on the position of the mapped points. Also, a local projection (LP) method is applied to refine the remaining ambiguous measures and increasing the quality of the reconstructed haplotypes.

The significant contributions of the proposed method are as follows:

- The similarity measurement between the input fragments can be described more accurately by utilizing the hypergraph model. Moreover, it helps to overcome challenges originated from the huge amount of gaps and sequencing errors.
- The quality score for each position of the reconstructed haplotypes can be calculated to predict the remaining error measures.
- The chaotic nature hypothesis is used to refine the reconstructed haplotypes. To this end, we only concentrate on the neighboring dependencies between SNPs.
- The proposed method could be applied effectively for both diploid and polyploid organisms.

The rest of the paper is organized as follows. Section 2 provides a brief review of the problem statement. In section 3, the proposed method is described in detail. Experimental results are presented in section 4. Finally, the conclusion is arrived at section 5.

## Preliminaries and assumptions

The challenge of the SIH problem in the polyploid organisms includes the reconstruction of the whole set *H* = {*h*_1_, *h*_2_, …, *h*_3_} containing P haplotype sequences. It is based on the available aligned input fragments. Similar to diploid case, the input fragments can be represented as a standard form. Let *X* be the SNP matrix in which each row corresponds to an input fragment, and each column indicates a specified SNP. In binary allelic haplotypes, it is assumed that *x*_*ij*_ ∈ {0,1,′ −′} indicating the obtained allele in a specified fragment *f*_*i*_ at SNP *s*_*j*_. Also, each haplotype *h*_*i*_ (*i* = 1,2, … , *P*) equals to {1,0}^N^. In diploid case, there are some positions called homozygote sites in which *h*_1*k*_ equals to *h*_2*k*_. On the other hand, the sites with different measures are called heterozygote positions. Homozygote sites are usually removed from the input matrix, as they do not provide useful information for the haplotype assembly problem. It is worth noting that the ′−′ sign indicates missing information during the sequencing process. For two fragments which are originated from different haplotypes, it is expected that there are some dissimilarities between them. Several relations have been developed to describe the differences between the two fragments. Hamming distance (HD) is the most practical approach, which can be used to calculate the differences between two input fragments *f*_*i*_ and *f*_*j*_ as follows:

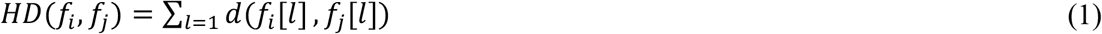

Where *d* is defined as follows:

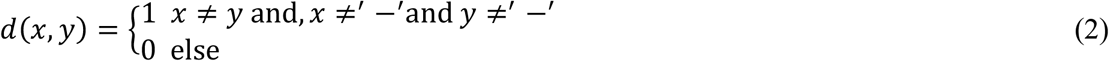

In the case where the SNP matrix is error-free, two fragments that were sequenced from the same haplotype are compatible, as their distance equals to zero. On the other hand, in dealing with the noisy SNP matrix, for two arbitrary fragments *f*_*i*_, *f*_*j*_, it is not possible to simply interpret the dissimilarity between two fragments, as they can be originated from the existing noise or have been sequenced from different haplotypes. In the error-free case, the fragments can be clustered in P clusters, such that the members of each cluster are compatible with each other.

Fig. 1 represents an example of the SIH problem in the ploidy level. The rows of matrix *X* indicate sequenced fragments, and the rows of matrix *H* contain the obtained haplotypes.

**Figure 1.**
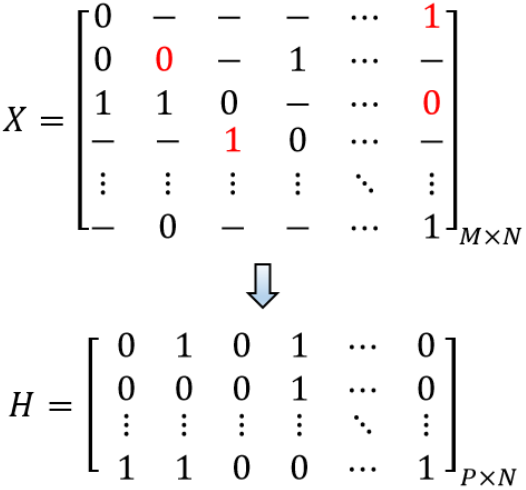
An example of SNP matrices *X* and *H* relevant to the resulting haplotypes. The red measures in *X* indicate sequencing errors. Each row of *H* demonstrates a specified haplotype sequence.

In diploid case, several models have been proposed to solve the SIH problem based on the input fragments.

Extending the models to solve the SIH problem in polyploidy form is a difficult task [3 8]. Recently, several MEC-based approaches have been developed to solve this problem. In this regard, the input fragments are organized in P clusters, and the haplotypes are considered as the centers of constructed clusters. In fact, each cluster involves the fragments which have the same provenance. The optimized result of the clustering algorithm can be obtained by minimizing the following Eq.:

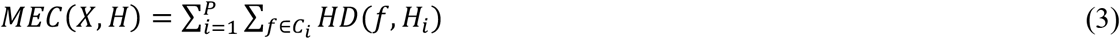

In the optimal case, if the SNP matrix is error-free, then the MEC measurement equals zero, and each fragment *f* belonging to *C*_*i*_ is compatible with *H*_*i*_. However, in dealing with the noisy SNP matrix, it is expected that some fragments to be in conflict with their corresponding haplotypes. It should be noted that finding the optimal MEC measure is an NP-hard problem. On the other hand, the huge amount of gaps in the input fragments does negatively affect the distance measurement between pairs of input fragments. Therefore, the current work aims to address these challenges by a better description of the similarity measurement between the input fragments. This was done by a heuristic method with a favorable runtime based on the hypergraph model.

## The proposed method

This section presents a <u>H</u>aplotype <u>R</u>econstruction approach based on the <u>C</u>haotic viewpoint and <u>H</u>ypergraph model (HRCH). The proposed method is briefly described below.

(i) a set of haplotype sequences is randomly generated;(ii) the input fragments are assigned to the haplotype sequences based on their similarities;(iii) a weighted SNP hypergraph is built, using the similarity measure between haplotype sequences and the assigned input fragments;(iv) the constructed hypergraph is used to find a set called CutSet, containing the SNPs which should be modified. This procedure is repeated for a predefined number of iterations to minimize the MEC score. Next, by considering the existence of chaotic properties of haplotype sequences, the results are improved. A high-level overview of the method is demonstrated in Fig. 2.

**Figure 2.**
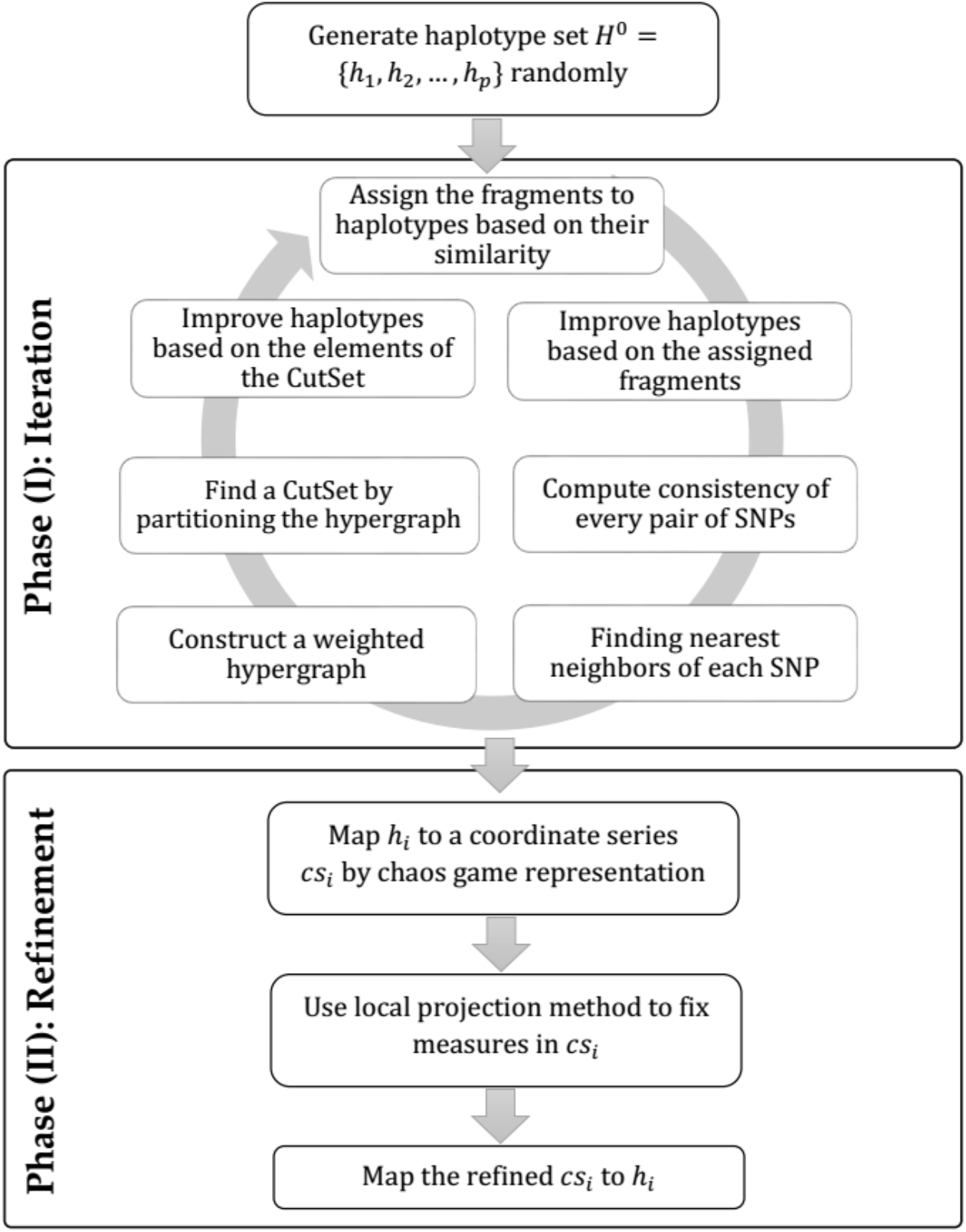
The workflow of proposed method.

### Data preprocessing

As described in the preliminaries sections, *X*_*M*×*N*_ is a matrix containing *M* reads with length *N*. It is essential to note that homozygote columns can be ignored in diploid cases. Removing the homozygote positions was performed as described by [33] such that the most frequent measure for each column could be found. If the frequency is higher than 0.8, the column is identified as a homozygote site. Thus, the output of this step is a matrix with *M* fragments and *N*′columns; where *N*′ ≤ *N*. Finally, *H*^0^ = {*h*_1_, *h*_2_, …, *h*_*p*_}, as an initial set of haplotypes is randomly generated.

### Pair-SNP consistency

Let ⋈ be a binary operator which provides the concatenation of two variables. For example, if *a* and *b* are two variables with measures ‘0’ and ‘1’, respectively, *a* ⋈ *b* equals to ‘01’. Given two variables *c*, *d* ∈ {′00′,′ 01′,′ 10′, ′11′}, the operator ⨁ is defined as follows:

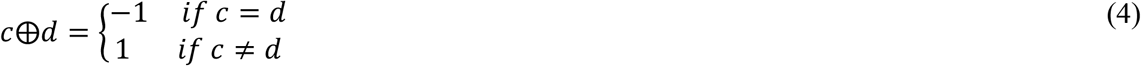

Definition 1 (Pair-SNP consistency). Given matrix *X*_*M*×*N*_ involving the input fragments, pair-SNP consistency, *ω*_*ij*_ is defined between *s*_*i*_ and *s*_*j*_ as two arbitrary SNPs which are covered by *f*_*k*_ *(k* = 1,2, …, *M)* as follows.

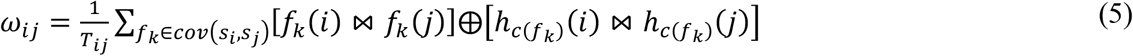

Where *T*_*ij*_ is the number of fragments covering both SNPs *s*_*i*_ and *s*_*j*_. By applying this measure, *ω*_*ij*_ is normalized such that its value ranges between −1 and +1 (*i.e.,* −1 ≤ *ω*_*ij*_ ≤ +1). Moreover, *cov*(*s*_*i*_, *s*_*j*_) includes the fragments which cover the SNPs *s*_*i*_ and *s*_*j*_. Finally, as mentioned above, *c*(*f*_*k*_) identifies the origin of *f*_*k*_.

The Pair-SNP consistency metric is used to evaluate the compatibility between each pair of SNPs with the current haplotype *H*^*t*^. The intuition behind the Eq. 5 is as follows. For given SNPs *s*_*i*_ and *s*_*j*_, *ω*_*ij*_ describes the amount of similarity between the pair SNPs and their corresponding measures in *H*^*t*^. This measure equals to −1 if the current haplotype is entirely identical with the covered fragments in columns *i* and *j*. On the other hand, it takes 1 if they are completely different in those columns. Otherwise, *ω*_*ij*_equals to 0, when the SNPs are not covered by any fragment. It is noticeable that, for high measures of *ω*_*ij*_, it is expected that the SNPs, *s*_*i*_, and *s*_*j*_, are considered to belong to different clusters upon partitioning. The complexity of this step is *O*(*MN*^2^), where *M* and *N* are the number of fragments and SNPs, respectively.

### Hypergraph construction

To construct the weighted hypergraph based on the achieved *ω* matrix, for each SNP *s*_*i*_, its K nearest neighbors is found using the following Eq.:

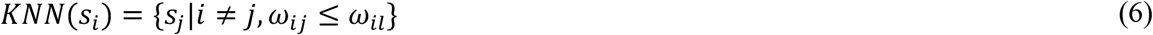

Where *l* is index of the *K*^*th*^ nearest SNP of *s*_*i*_. *KNN*(*s*_*i*_) is a set containing the index of *K* nearest neighbors of *s*_*i*_. More specifically, each set represents the *K* SNPs, which have the most consistent relationship with *s*_*i*_. In this case, each hyperedge can connect more than two vertices. Applying K nearest neighbors is a common approach to determine the hyperedges. However, it is necessary to specify the hyperedges more precisely due to the existing noise and sparsity of the SNP matrix. Therefore, the connectivity of vertices is defined by finding frequent itemsets. In other words, the hyperedges are determined as the shared K nearest neighbors, which can be defined as follows:

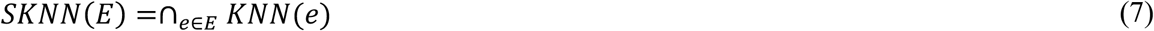

In Eq. 7, *E* contains several SNPs, and *SKNN*(*E*) provides a set of SNPs which are shared between all nearest neighbors of *E*. If the number of shared KNNs is more than a predefined threshold, called minimum support count (sc), then E can be defined as a frequent itemset. In the proposed model, each frequent itemset is defined as a specified hyperedge *e*_*i*_, and the number of shared KNN is assigned as its weight measure *w*_*i*_. Among the existing methods, frequent pattern (FP)-growth [39] has been gaining much attention due to the ability to find frequent itemsets. FP-growth is a tree-based method which uses a depth-first strategy to mine frequent itemsets. Accordingly, the database is modeled as a prefix tree, and the depth-first search is recursively applied to generate all maximal frequent itemsets. The runtime of this algorithm increases linearly, and it depends on the number of SNPs[40].

### Improving *H*^*t*^ by partitioning the hypergraph

As can be seen in Fig. 3, in the constructed hypergraph, the SNPs correspond with vertices, and each hyperedge equals with an obtained frequent itemset. In other words, it contains a set of SNPs that has more consistency with the corresponding position in *H*^*t*^. It is noteworthy that hyperedges with higher weights indicate the higher similarity between the constituent’s SNPs and their relevant positions in *H*^*t*^. The vertices can be divided into two clusters via partitioning the hypergraph. The objective of the partitioning is to minimize the sum of the weights of the hyperedges located between the clusters. To this end, hmetis as a popular algorithm was used. The algorithm includes three steps: (i) a number of small hypergraphs in several layers are built; (ii) the hypergraph in the lowest level is partitioned; (iii) the resulted partitions are extended to the upper levels through a successive mapping.

**Figure 3.**
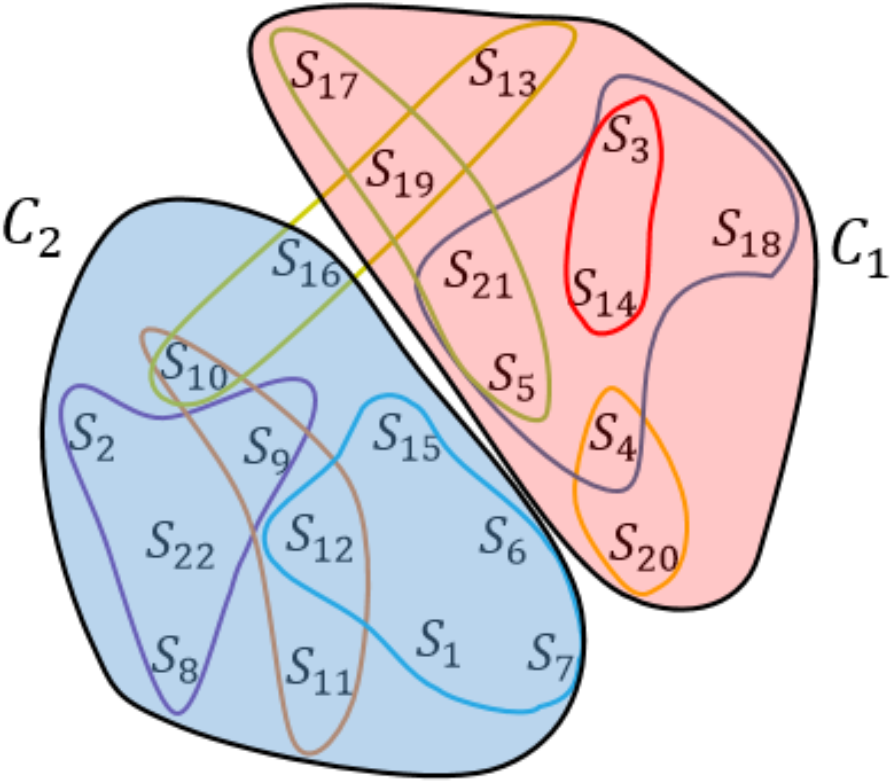
An example of constructing and partitioning the hypergraph. *S*_*i*_ corresponds with the i^th^ SNP, and the curves demonstrate the hyperedges. *C*_1_ and *C*_2_ denote the clusters which are obtained by hypergraph partitioning.

The computational complexity of the algorithm is *O*(|*E*|), where *E* is the set of hyperedges. Suppose that *C*_1_ and *C*_2_ are two clusters obtained by the hmetis algorithm. As can be seen in Fig. 4, 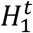 and 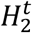 are partial haplotypes originating from the resulting clusters.

**Figure 4.**
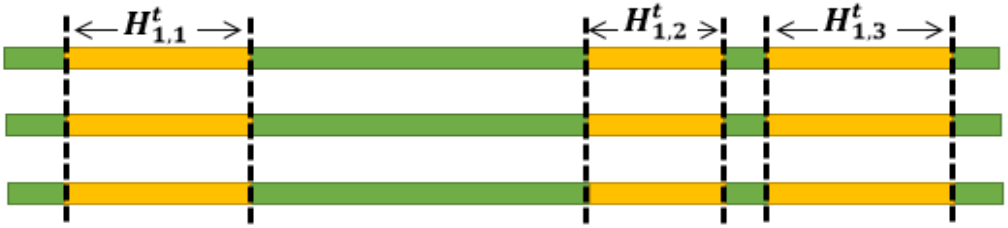
Partitioning *H*^*t*^ of a three ploid genome. The yellow parts indicate 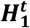 and the green parts demonstrate 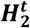. It must be pointed out that 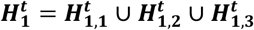.

In the diploid case, as can be seen in Fig.5 like the HapCUT method, improving *H*^*t*^ is performed as follows. First, *C*_1_ or *C*_2_ is selected as a CutSet. Next, *H*^t+1^ is obtained from *H*^*t*^ by flipping the measures of the SNPs in the CutSet.

**Figure 5.**
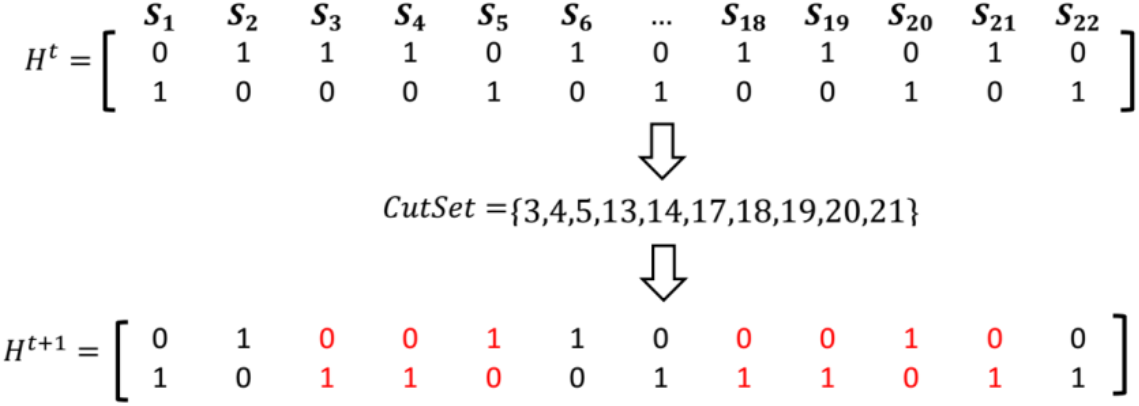
An example of updating the current haplotype based on the partitioning of the hypergraph.

For polyploid, improving *H*^*t*^ is accomplished based on the algorithm which is shown in Fig. 6. In the first step, a partial haplotype (i.e., 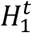 or 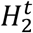) is randomly assigned to CutSet. This set involves some parts of *H*^*t*^ that should be corrected.

**Figure 6.**
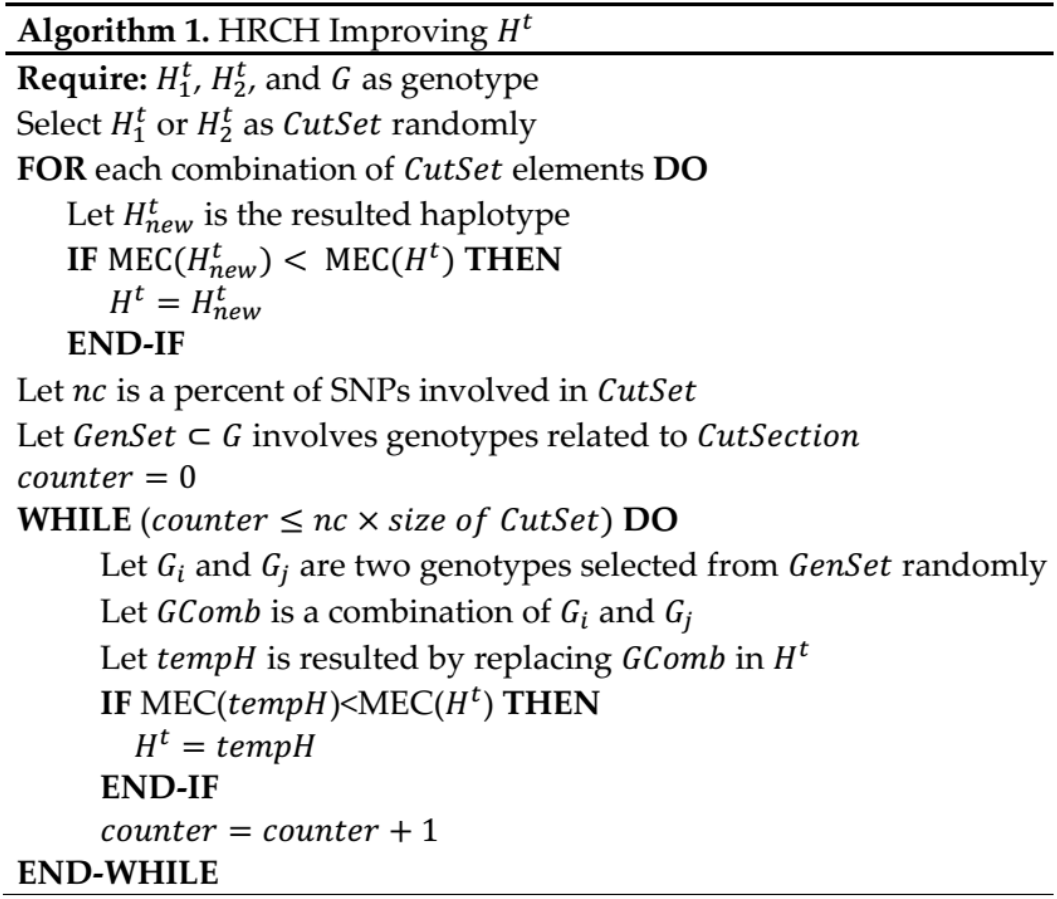
The algorithm of improving *H*^*t*^.

As shown in Fig. 7, all combinations of the CutSet are evaluated to find a new set of haplotypes 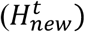 with lower MEC score.

**Figure 7.**
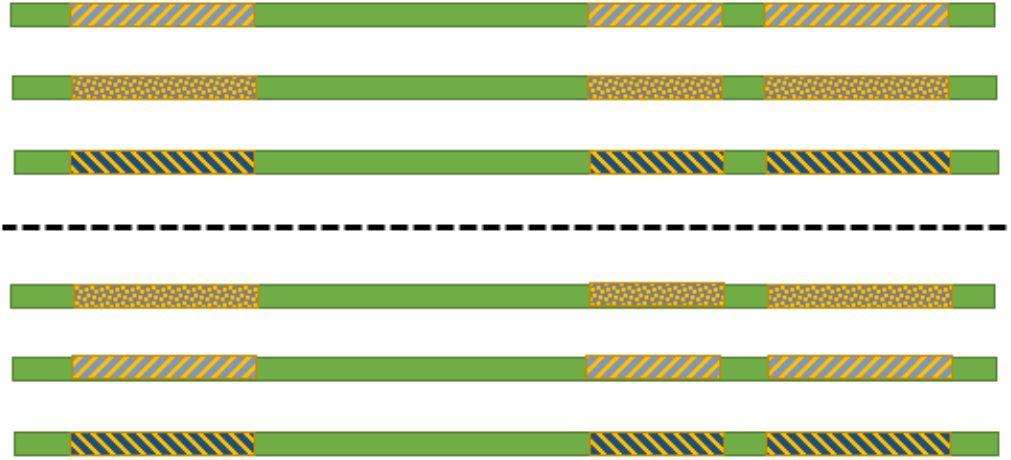
Two combinations of six possible combinations of the CutSet in three ploid form.

Moreover, in order to evaluate more allelic combinations of SNPs, for a predefined percent of SNPs belonging to the CutSet, in each time two arbitrary SNPs are nominated. Then one of its various genotype’s combinations is randomly selected, and is replaced at corresponding positions in *H*^*t*^. This step repeats for a predefined percent of SNPs.

Since *H*^*t*^ has randomly generated, in the early iterations, its MEC score is poor. Therefore, finding the hyperedges with lower weights is not a difficult task. But, by improving the quality of *H*^*t*^ and increasing the consistencies between SNPs, MEC measure will be decreased slowly.

## Refinement of *H*^*t*^

### Computing confidence score

Upon performing the iterative procedure of the proposed method, the haplotype *H* = {*h*_1_, *h*_2_, …, *h*_*p*_} will be obtained. It is possible to define a confidence measure for each loci of the reconstructed haplotypes. For diploid case, we used the emission probability *P*(*X*_*j*_|*h*_*j*_, *R*_*j*_) that has been defined in [41], which is used to identify errors in the reconstructed haplotype. This measure is calculated for each position *j* as follows:

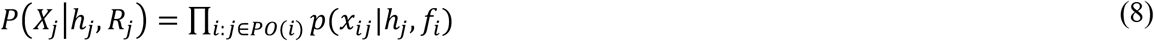

Where *h* is a haplotype sequence belongs to *H*, *h*_*j*_ ∈ {0,1} denotes an allele in position *j*, and *PO*(*i*) contains the columns which have been covered by *f*_*i*_ as the i-th fragment. Furthermore, *R*_*j*_ is a set which includes fragments such as *f*_*i*_ covering a position *j*. Finally, *p*(*x*_*ij*_|*h*_*j*_, *f*_*i*_) is calculated as follows:

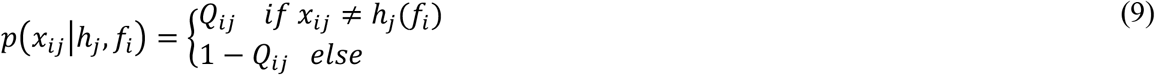

Where *Q* is an *M* × *N* matrix; for each element *x*_*ij*_ ∈ *X*, *Q*_*ij*_ includes the probability of sequencing error and *h*_*j*_(*f*_*i*_) as the j-th loci of the reconstructed haplotype is computed based on the following Eq.:

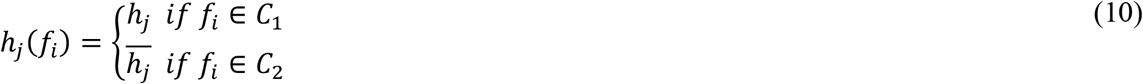

Where *C*_1_ and *C*_2_ are the obtained clusters containing similar fragments that indicate the provenance of *f*_*i*_. Eq. (10) provides more information in each loci, and is based on the fact that 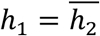. Therefore, the confidence score could be calculated more precisely. On the other hand, there is no relationship between the haplotype sequences in the polyploid form. Hence, applying Eq. (8) is not applicable. In this case, we used genotype information. Suppose that *g*_*i*_ is the genotype information in position *i* and *H*_*i*_ is the reconstructed measure in this position. The sorted measure of *g*_*i*_ and *H*_*i*_ are compared, and the position i will be selected for refinement if the two sets are not equivalent.

### Applying chaos game representation

Chaos game representation (CGR) is a graphical tool which maps an arbitrary sequence to a 2-dimensional form.

This map is reversible and all the information of the sequence is preserved. Moreover, it depicts the hidden dependencies among the letters. CGR was initially introduced by Barnsley[42] to evaluate random sequences. Afterwards, Jeffrey[43] developed the method for visualizing genomic sequences. For this aim, according to the number of distinct letters constructing the input sequence, a regular polygon can be considered. For example since DNA sequences are constructed from four nucleotides ‘a’, ‘t’, ‘c’, and ‘g’, a square with unit length is considered and each distinct letter is assigned to one vertex. Each letter of the given sequence is iteratively mapped as a point inside the square. The process is started by locating the first point half-way between the center of the square and the corner related to the occurrence of the first letter. The method continues such that the i-th point is placed half-way between the previous point and the vertex related to the i-th letter. Using this procedure, many attempts have been made with the purpose of extracting novel features from biological sequences by exploiting CGR[44–48].

Recently, CGR was used to reveal the chaotic properties of haplotypes[37]. Since haplotypes are represented in binary form, the achieved map will be a dotted line which its vertices are named by 0 and 1, respectively. In this step, in order to improve the reconstructed haplotypes, CGR is utilized as follows.

For loci’s which their qualities are less than *θ*, as a predefined threshold, their measures may be disrupted by noise or missing information. Therefore, it is refined based on the existing dependencies between SNPs. For this purpose each *h*_*i*_ ∈ *H* is mapped to a line by applying CGR. The places assigned to each point construct a coordinate series, namely *cs*_*i*_. The route of chaos helps to refine the ambiguous measures in the low-quality positions. To this end, the points in *cs*_*i*_ that are correspond with the alleles with low confidences, are shown by ‘-‘. Then, the measure of ambiguous positions can be determined by applying a local projection (LP) method. After filling the removed measures based on the LP method, the refined coordinate series called 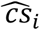, are transformed into the final haplotype known as 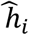. It must be indicated that extracting the *cs* and applying the LP method are accomplished in linear time. The conversion is calculated according to Eq. 11:

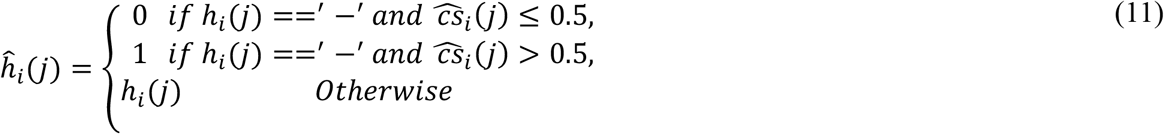

## Results

In the following section, the performance of the proposed method is compared with several state-of-the-art approaches in diploid and polyploid forms. The method was implemented in MATLAB, and all the results were obtained on a Windows 10 PC with 3.6 GHz CPU and 16 G Ram. The parameters of the algorithm are defined as *t* = 100, *k* = 5, *sc* = 2, and *nc* = 20%. Reconstruction rate (RR)[4] as a conventional metric was used to evaluate the quality of the obtained haplotypes. In diploid case, RR is defined as follows:

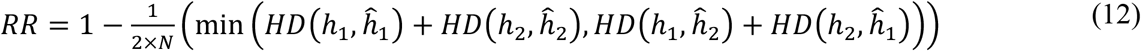

Here, *HD* denotes hamming distance between *h*_*i*_ and 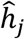 which are the target and the reconstructed haplotype, respectively and *i, j* = 1,2. For polyploid case, this formula is written in the form of Eq. 13:

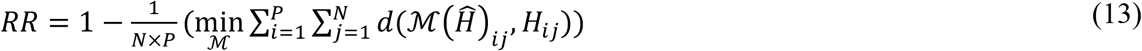

Where ℳ is a one-to-one mapping from the set of reconstructed haplotypes to the set of target haplotypes.

### Diploid case

The experiments have been carried out on two widely used and well-known datasets including Geraci’s dataset[49] and a dataset from the 1000 genome project that are prime examples of the simulated and experimental datasets, respectively.

### Simulated data

The Geraci’s dataset involves three parameters: Coverage *c* = {3,5,8,10}, length of haplotypes *l* = {100,350,700}, and error rate *e* = {0,0.1,0.2,0.3}. For each combination of these parameters, there are 100 samples. The output of the proposed method was compared with a set of state-of-the-art and well-known methods including; SCGD[36], H-pop[34], ARO[24], HG[33], FCM[25], FastHap[26], DGS[50], SHR[51], MLF[52], HapCut[27], 2d[22], Fast[53], and SPH[54]. All of these methods were run with their default parameter settings. In accordance with the existing methods, the reconstruction rate (RR) was also used to assess the result of the current method. Tables 1, 2 and, 3 comparatively show the reconstruction rate of the proposed method with those described for haplotype blocks with length 100, 350, and 700. Note that in the last column of each table, the highest measures are boldfaced. Moreover, the second-highest measures are highlighted. The results, demonstrate that current method has an acceptable level of performance and outperforms in most of the cases.

**Table 1.**
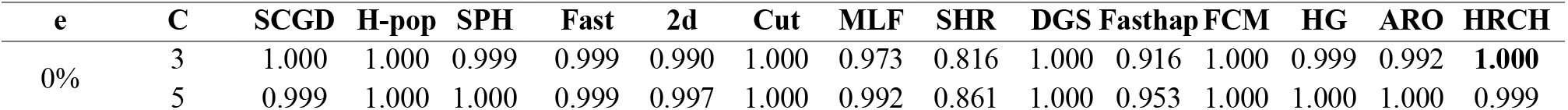
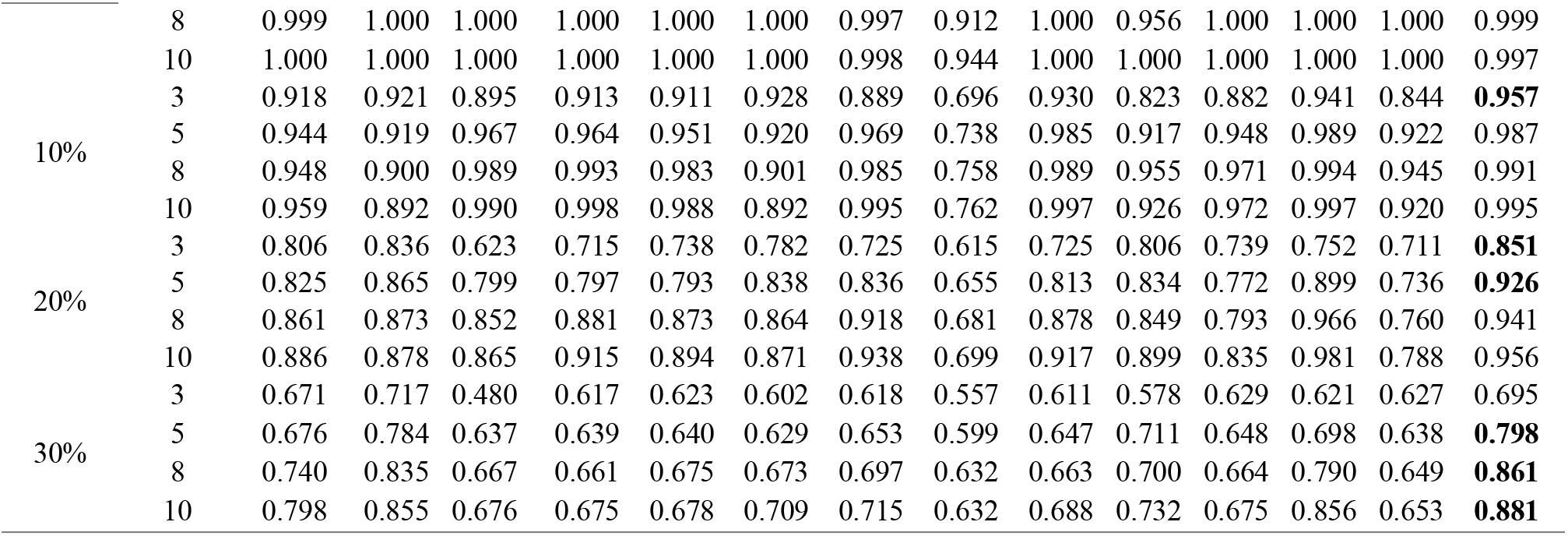
Average of reconstruction rate for haplotypes with length 100.

**Table 2.**
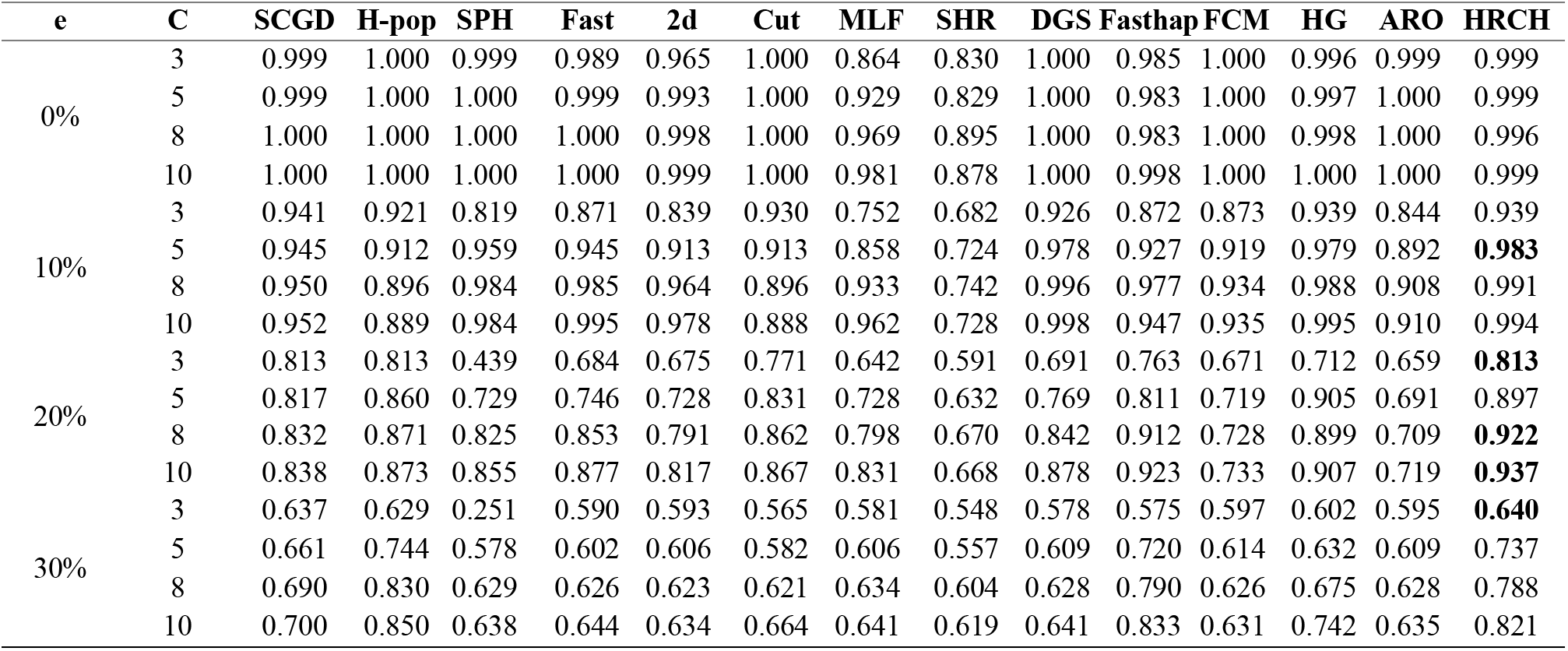
Average of reconstruction rate for haplotypes with length 350.

**Table 3.**
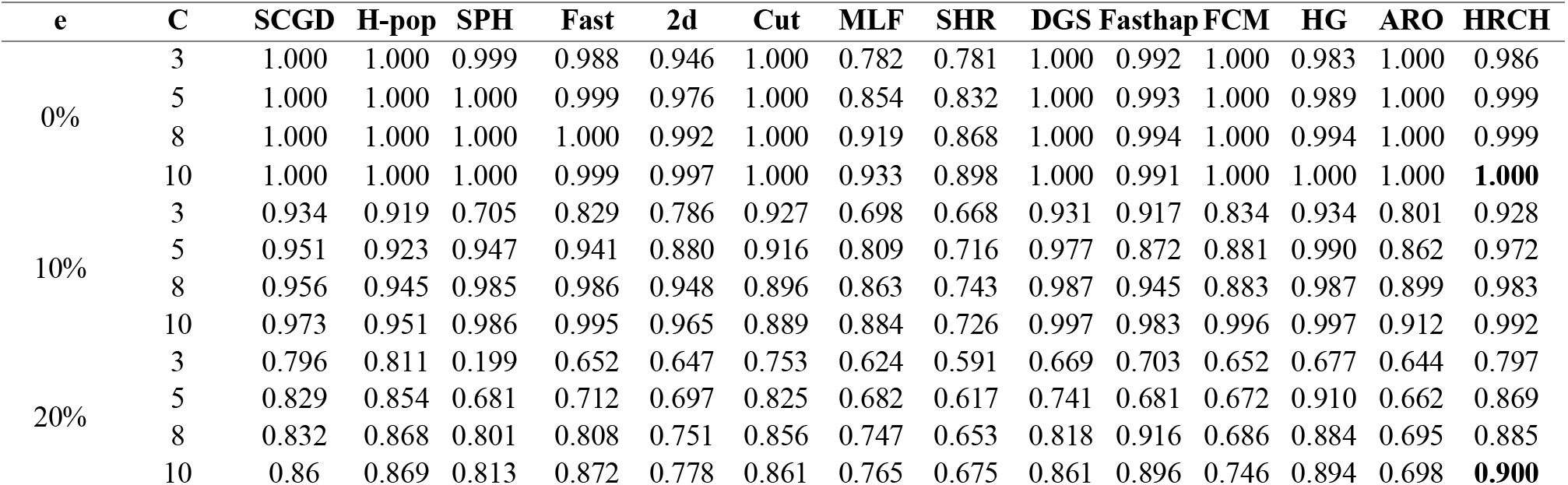
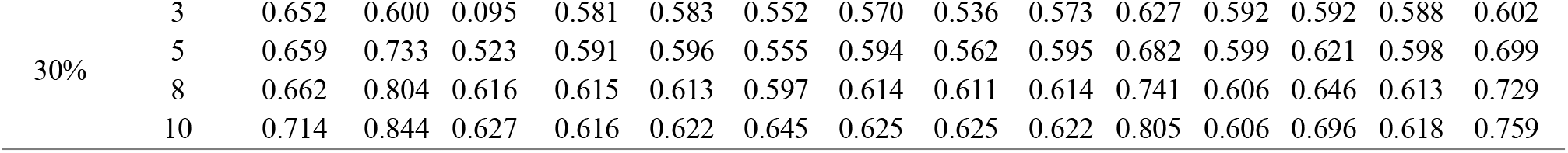
Average of reconstruction rate for haplotypes with length 700.

The performance of the refinement phase has been considered in Table 4. Since evaluating the chaotic feature is limited to the long coordinate series, this phase can only be performed for sequences with length 700. For this purpose, the LP method is applied for each coordinate series with embedding dimensions (*em*) equal to 1 and 2, individually. It should be noted that the first column demonstrates the quality of the obtained haplotypes after terminating the first phase. The next two columns involve the rate of reconstruction for em equals to 1 and 2, respectively. The obtained results demonstrate that the inclusion of the chaotic nature of haplotype sequences can significantly improve the reconstruction rate.

**Table 4.** The effect of refinement phase for haplotypes with length 700 in diploid case.

| E | C | Hypergraph | em=1 | em=2 |
| --- | --- | --- | --- | --- |
| 10% | 3 | 0.916 | 0.918 | 0.928 |
|  | 5 | 0.966 | 0.968 | 0.972 |
|  | 8 | 0.980 | 0.981 | 0.983 |
|  | 10 | 0.989 | 0.991 | 0.992 |
| 20% | 3 | 0.786 | 0.787 | 0.797 |
|  | 5 | 0.854 | 0.857 | 0.869 |
|  | 8 | 0.870 | 0.880 | 0.885 |
|  | 10 | 0.886 | 0.896 | 0.900 |
| 30% | 3 | 0.600 | 0.601 | 0.602 |
|  | 5 | 0.688 | 0.689 | 0.699 |
|  | 8 | 0.716 | 0.720 | 0.729 |
|  | 10 | 0.748 | 0.754 | 0.759 |

### Experimental dataset

The second dataset which is used for evaluation of the proposed algorithm involves experimental data which was provided by 1000 genome project. The gathered data belongs to an individual NA12878 which often is used to analyze the performance of the existing haplotype assembly methods. The sample was provided by using 454 sequencing method. According to the overlapping of the obtained fragments, they are represented in multiple matrices. In this experiment, for each chromosome, the first 500 matrices have been selected. In each matrix, the length of each row is ~90 in average and cover the genome at a depth of ~ × **3**. Furthermore, the trio-phased variant calls from the GATK resource bundle[55] was used as the target haplotypes. The obtained reconstruction rates of the proposed method are compared to those of H-pop[34], SCGD[36], HG[33], ARO[24], and FCM[25] approaches. The results for all 22 homologous chromosomes are listed in Table 5. The results show that in most cases the proposed method achieved higher reconstruction rates compared to the others. The last row of the table demonstrates the mean of RR values of the comparing methods for all of the chromosomes. According to the obtained results, it can be concluded that this method completely outperforms the other approaches.

**Table 5.** The reconstruction rate and running time for the proposed method, H-pop, SCGD, HG, ARO and FCM applied to the experimental dataset NA12878 dataset provided by 1000 genome project.

| Chr | H-pop |  | SCGD |  | HG |  | ARO |  | FCM |  | HRCH |  |
| --- | --- | --- | --- | --- | --- | --- | --- | --- | --- | --- | --- | --- |
|  | RR | t(sec) | RR | t(sec) | RR | t(sec) | RR | t(sec) | RR | t(sec) | RR | t(sec) |
| 1 | 0.957 | 5.22 | 0.925 | 3.62 | 0.937 | 1.54 | 0.935 | 20.28 | 0.913 | 1.09 | 0.954 | 10.40 |
| 2 | 0.956 | 5.65 | 0.926 | 4.41 | 0.929 | 1.30 | 0.943 | 18.03 | 0.908 | 1.04 | 0.943 | 12.34 |
| 3 | 0.912 | 6.99 | 0.919 | 3.40 | 0.928 | 1.17 | 0.940 | 18.45 | 0.913 | 1.91 | **0.944** | 12.75 |
| 4 | 0.970 | 5.24 | 0.927 | 5.47 | 0.923 | 1.20 | 0.949 | 18.06 | 0.923 | 1.68 | 0.960 | 13.07 |
| 5 | 0.966 | 4.67 | 0.939 | 3.54 | 0.932 | 1.24 | 0.942 | 15.09 | 0.912 | 1.27 | 0.952 | 14.98 |
| 6 | 0.952 | 4.93 | 0.930 | 8.70 | 0.935 | 1.22 | 0.948 | 15.60 | 0.929 | 1.04 | **0.958** | 13.58 |
| 7 | 0.924 | 4.24 | 0.935 | 3.95 | 0.925 | 1.26 | 0.951 | 16.34 | 0.904 | 1.03 | **0.954** | 12.53 |
| 8 | 0.947 | 4.14 | 0.907 | 2.18 | 0.906 | 1.25 | 0.934 | 16.62 | 0.903 | 1.07 | **0.949** | 13.03 |
| 9 | 0.910 | 3.36 | 0.971 | 2.94 | 0.901 | 1.30 | 0.966 | 15.25 | 0.937 | 1.04 | 0.921 | 12.63 |
| 10 | 0.945 | 3.67 | 0.926 | 2.56 | 0.940 | 1.21 | 0.945 | 15.73 | 0.913 | 1.28 | **0.954** | 13.14 |
| 11 | 0.915 | 3.71 | 0.932 | 2.95 | 0.939 | 1.17 | 0.942 | 14.34 | 0.923 | 1.18 | **0.963** | 10.46 |
| 12 | 0.903 | 3.46 | 0.923 | 2.03 | 0.945 | 1.19 | 0.935 | 14.26 | 0.908 | 1.14 | **0.954** | 11.33 |
| 13 | 0.941 | 2.89 | 0.970 | 3.31 | 0.930 | 1.22 | 0.935 | 15.72 | 0.925 | 1.43 | 0.946 | 14.12 |
| 14 | 0.971 | 2.54 | 0.911 | 1.36 | 0.917 | 1.52 | 0.934 | 15.42 | 0.932 | 1.11 | 0.949 | 14.03 |
| 15 | 0.974 | 2.40 | 0.991 | 1.21 | 0.920 | 1.02 | 0.937 | 16.65 | 0.905 | 1.04 | 0.951 | 12.24 |
| 16 | 0.935 | 2.47 | 0.930 | 1.79 | 0.932 | 1.11 | 0.946 | 15.27 | 0.924 | 1.35 | **0.962** | 11.01 |
| 17 | 0.911 | 1.98 | 0.967 | 2.61 | 0.931 | 1.25 | 0.951 | 15.86 | 0.920 | 1.11 | 0.963 | 11.35 |
| 18 | 0.976 | 2.51 | 0.903 | 1.16 | 0.924 | 1.86 | 0.949 | 15.66 | 0.919 | 1.01 | 0.954 | 13.02 |
| 19 | 0.978 | 1.82 | 0.972 | 3.25 | 0.949 | 1.60 | 0.942 | 14.58 | 0.923 | 1.40 | 0.960 | 10.46 |
| 20 | 0.950 | 2.00 | 0.968 | 1.38 | 0.945 | 1.90 | 0.946 | 15.49 | 0.922 | 1.12 | **0.957** | 11.31 |
| 21 | 0.970 | 1.70 | 0.943 | 0.63 | 0.933 | 1.52 | 0.941 | 15.12 | 0.915 | 1.08 | 0.960 | 12.77 |
| 22 | 0.983 | 1.44 | 0.941 | 0.74 | 0.951 | 1.16 | 0.941 | 14.34 | 0.914 | 1.33 | 0.964 | 9.64 |
| Mean | 0.948 | 3.50 | 0.939 | 2.87 | 0.931 | 1.33 | 0.943 | 16.00 | 0.918 | 1.22 | **0.953** | 12.28 |

### Polyploid case

Here, the proposed method is compared with three recent approaches that have been developed to solve haplotype assembly in polyploid form including Althap[23], H-POP[34] and SCGD[36]. The source codes of all comparing methods are available. To investigate the quality of reconstructed haplotypes, reconstruction rate (RR), and MEC measure of the methods have compared.

Indeed, the benchmark dataset is provided by simulation. For this aim, we have used the source code, which is available upon request by [23]. Its input parameters are coverage (*c*), error rate (*e*), and length of haplotypes (*l*). In this experiment, we defined *c* ∈ {5,10,15,20}, *e* ∈ {0.1,0.2,0.3} and *l* ∈ {100,350,700}. For each combination of those parameters 10 samples have generated. Each sample contains an SNP matrix with a huge amount of gaps. As can be seen in Tables 6, 7, and 8 the proposed method is compared with RR and MEC-based algorithms.

**Table 6.** Average of reconstruction rate for haplotypes with length 100.

| e | C | SCGD |  | H-pop |  | AltHap |  | HRCH |  |
| --- | --- | --- | --- | --- | --- | --- | --- | --- | --- |
|  |  | RR | MEC | RR | MEC | RR | MEC | RR | MEC |
| 10% | 5 | 0.609 | 1289 | 0.745 | 269 | 0.736 | 315 | **0.830** | **260** |
|  | 10 | 0.567 | 2917 | 0.813 | 534 | 0.783 | 598 | **0.846** | **523** |
|  | 15 | 0.567 | 4282 | 0.828 | 805 | 0.754 | 1095 | **0.859** | **773** |
|  | 20 | 0.564 | 5846 | **0.844** | **1004** | 0.747 | 1864 | 0.839 | 1020 |
| 20% | 5 | 0.596 | 1367 | 0.667 | 467 | 0.657 | 478 | **0.717** | **447** |
|  | 10 | 0.548 | 2862 | 0.692 | 1009 | 0.730 | 1050 | **0.768** | **942** |
|  | 15 | 0.549 | 3047 | 0.706 | 1539 | 0.666 | 2241 | **0.797** | **1443** |
|  | 20 | 0.547 | 5894 | 0.740 | 2016 | 0.631 | 3559 | **0.781** | **1920** |
| 30% | 5 | 0.587 | 1373 | 0.596 | 554 | 0.630 | 548 | **0.661** | **538** |
|  | 10 | 0.550 | 2857 | 0.599 | 1244 | 0.633 | 1338 | **0.689** | **1190** |
|  | 15 | 0.548 | 4267 | 0.619 | 1916 | 0.596 | 2640 | **0.730** | **1849** |
|  | 20 | 0.550 | 5916 | 0.633 | 2591 | 0.572 | 4051 | **0.730** | **2512** |

**Table 7.**
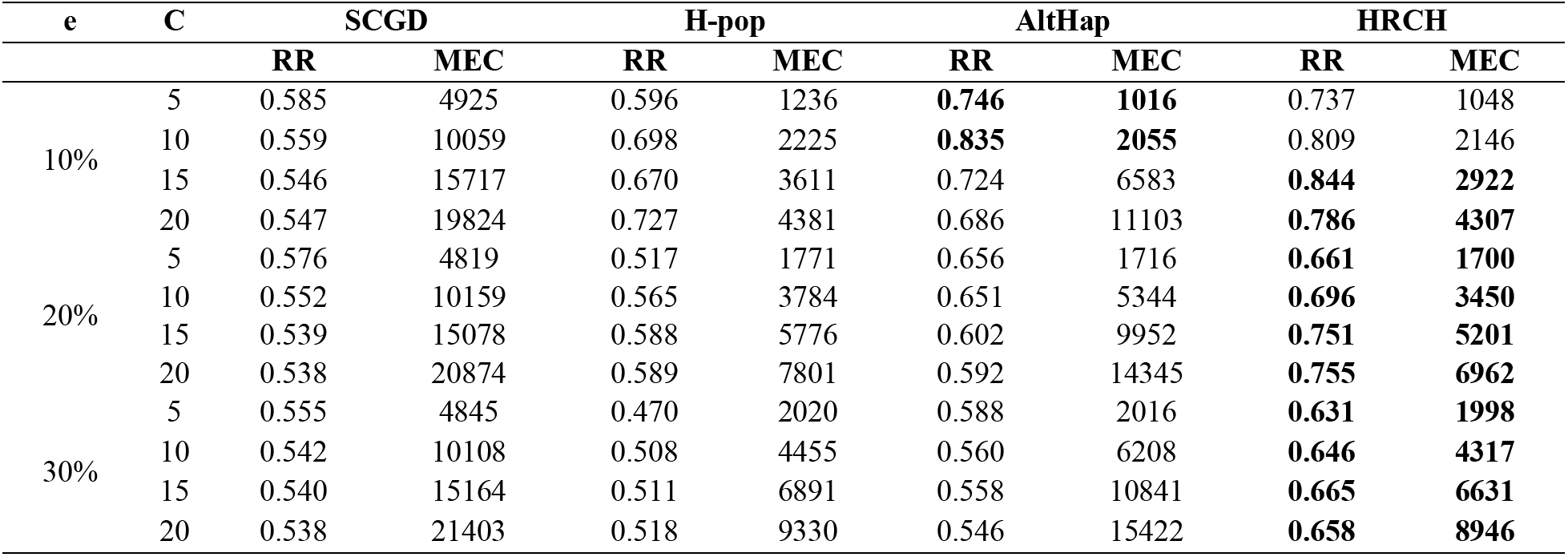
Average of reconstruction rate for haplotypes with length 350.

| e | C | SCGD |  | H-pop |  | AltHap |  | HRCH |  |
| --- | --- | --- | --- | --- | --- | --- | --- | --- | --- |
|  |  | RR | MEC | RR | MEC | RR | MEC | RR | MEC |
| 10% | 5 | 0.585 | 4925 | 0.596 | 1236 | **0.746** | **1016** | 0.737 | 1048 |
|  | 10 | 0.559 | 10059 | 0.698 | 2225 | **0.835** | **2055** | 0.809 | 2146 |
|  | 15 | 0.546 | 15717 | 0.670 | 3611 | 0.724 | 6583 | **0.844** | **2922** |
|  | 20 | 0.547 | 19824 | 0.727 | 4381 | 0.686 | 11103 | **0.786** | **4307** |
| 20% | 5 | 0.576 | 4819 | 0.517 | 1771 | 0.656 | 1716 | **0.661** | **1700** |
|  | 10 | 0.552 | 10159 | 0.565 | 3784 | 0.651 | 5344 | **0.696** | **3450** |
|  | 15 | 0.539 | 15078 | 0.588 | 5776 | 0.602 | 9952 | **0.751** | **5201** |
|  | 20 | 0.538 | 20874 | 0.589 | 7801 | 0.592 | 14345 | **0.755** | **6962** |
| 30% | 5 | 0.555 | 4845 | 0.470 | 2020 | 0.588 | 2016 | **0.631** | **1998** |
|  | 10 | 0.542 | 10108 | 0.508 | 4455 | 0.560 | 6208 | **0.646** | **4317** |
|  | 15 | 0.540 | 15164 | 0.511 | 6891 | 0.558 | 10841 | **0.665** | **6631** |
|  | 20 | 0.538 | 21403 | 0.518 | 9330 | 0.546 | 15422 | **0.658** | **8946** |

**Table 8.**
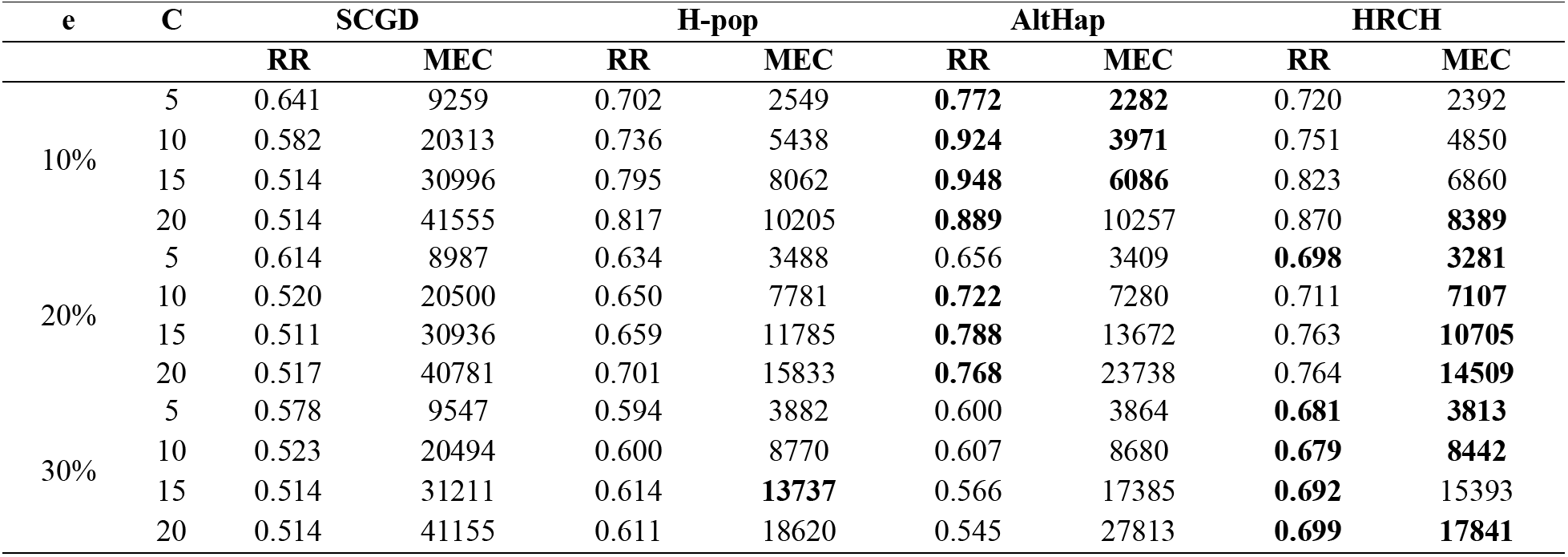
Average of reconstruction rate for haplotypes with length 700.

The results demonstrate that the proposed method outperforms the other approaches in most cases considering both RR and MEC parameters. Similar to the previous section, to emphasize the efficiency of the refinement phase, the RRs of haplotypes with length 700 have been considered, as provided in Table 9. Similar to the diploid case, the improvement of RRs reveals the role of chaotic viewpoint to efficiently decrease the amount of remaining noise in the constructed haplotypes. Obviously, the proposed method is slower than the competitors, because it starts from a random measure and is iterative. However, it can solve the problem in a reasonable amount of time.

**Table 9.** The effect of refinement phase for haplotypes with length 700 in polyploid case.

| e | C | Hypergraph | em=1 | em=2 |
| --- | --- | --- | --- | --- |
| 10% | 5 | 0.712 | 0.715 | 0.720 |
|  | 10 | 0.749 | 0.750 | 0.751 |
|  | 15 | 0.823 | 0.823 | 0.823 |
|  | 20 | 0.870 | 0.870 | 0.870 |
| 20% | 5 | 0.678 | 0.687 | 0.698 |
|  | 10 | 0.705 | 0.708 | 0.711 |
|  | 15 | 0.762 | 0.763 | 0.763 |
|  | 20 | 0.763 | 0.764 | 0.764 |
| 30% | 5 | 0.643 | 0.662 | 0.681 |
|  | 10 | 0.664 | 0.672 | 0.679 |
|  | 15 | 0.681 | 0.688 | 0.692 |
|  | 20 | 0.692 | 0.697 | 0.699 |

Since sequencing coverage of the used benchmark datasets were relatively low, we further evaluated the performance of HRCH by dealing with high coverage data. For this aim, by using the provided source code in [23], for diploid and polyploid cases, several samples were generated individually. For each combination of l=500, e=0.3, and c={30,40,50}, 10 samples were generated. As shown in Fig. 8, the reconstruction rates of the proposed method are compared to those of AltHap[23], SCGD[36], and H-pop[34]. The obtained results demonstrate that HRCH provides encouraging accuracy as compared to the competing schemes in diploid and polyploid forms.

**Figure 8.**
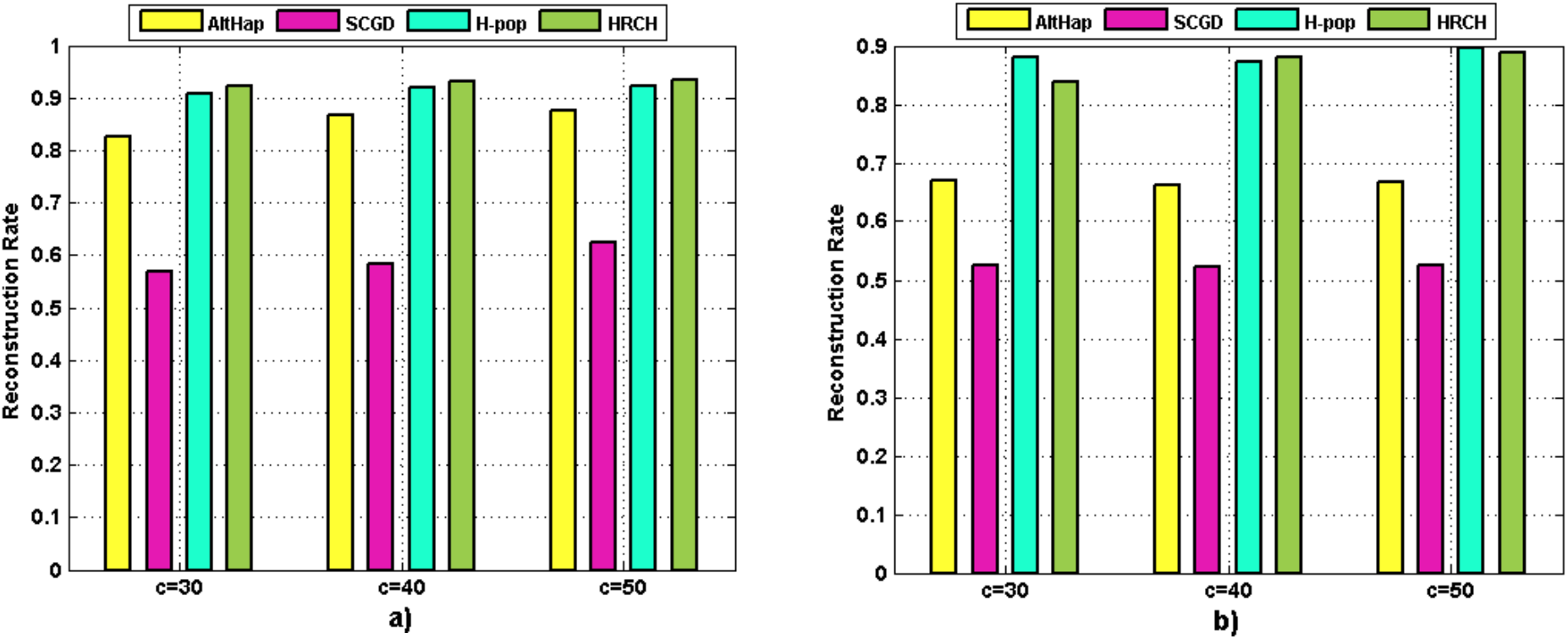
Comparison of reconstruction rate of the methods over high coverage data a) Diploid b) Polyploid.

## Conclusion

The high amounts of noise, as well as existing gaps in the input fragments, are the main challenges in solving the SIH problem. In this study, we established a sampling-based method that starts from an initial set of haplotypes and iteratively proceeds to improve the input data by correcting the SNPs with wrong measures. The proposed method involves two main steps. First, it utilizes the hypergraph model to conquer the sparsity and high amount of noise, and reconstructs haplotypes iteratively. Positions with low confidence are then rectified by mapping haplotype sequences to the coordinate series and applying a chaotic viewpoint. The proposed method has the capability to manipulate genomic data of both diploid and polyploid organisms. The promising results for diploid and polyploid data highlight that the method is comparable with the existing approaches, and they have complementary roles to each other. Finally, the source code of the proposed method is available at https://github.com/mholyaee/HRCH.

## Acknowledgments

The authors wish to thank Dr. Jamshid Pirgazi and Dr. Omid AbbasZadeh for their valuable suggestions and discussions. We also thank Dr. Sajad Ahmadian and Dr. Sina Majidian for their constructive comments.

## References

1. Browning SR, Browning BL (2011) Haplotype phasing: existing methods and new developments. Nature Reviews Genetics 12: 703.

2. Group ISMW (2001) A map of human genome sequence variation containing 1.42 million single nucleotide polymorphisms. Nature 409: 928.

3. Lee C-Y (2016) A model for the clustered distribution of SNPs in the human genome. Computational Biology and Chemistry 64: 94–98.

4. Wang R-S, Wu L-Y, Li Z-P, Zhang X-S (2005) Haplotype reconstruction from SNP fragments by minimum error correction. Bioinformatics 21: 2456–2462.

5. Loggetto SR (2013) Sickle cell anemia: clinical diversity and beta S-globin haplotypes. Revista brasileira de hematologia e hemoterapia 35: 155–157.

6. Rohlfs EM, Zhou Z, Heim RA, Nagan N, Rosenblum LS, et al. (2011) Cystic fibrosis carrier testing in an ethnically diverse US population. Clinical chemistry 57: 841–848.

7. McLaren GD, Gordeuk VR (2009) Hereditary hemochromatosis: insights from the hemochromatosis and iron overload screening (HEIRS) study. ASH Education Program Book 2009: 195–206.

8. Wilson JF, Weale ME, Smith AC, Gratrix F, Fletcher B, et al. (2001) Population genetic structure of variable drug response. Nature genetics 29: 265.

9. Exner DV, Dries DL, Domanski MJ, Cohn JN (2001) Lesser response to angiotensin-converting – enzyme inhibitor therapy in black as compared with white patients with left ventricular dysfunction. New England Journal of Medicine 344: 1351–1357.

10. Varner RV, Ruiz P, Small DR (1998) Black and white patients response to antidepressant treatment for major depression. Psychiatric Quarterly 69: 117–125.

11. Glusman G, Cox HC, Roach JC (2014) Whole-genome haplotyping approaches and genomic medicine. Genome medicine 6: 73.

12. Lawson DJ, Hellenthal G, Myers S, Falush D (2012) Inference of population structure using dense haplotype data. PLoS genetics 8: e1002453.

13. Green RE, Krause J, Briggs AW, Maricic T, Stenzel U, et al. (2010) A draft sequence of the Neandertal genome. science 328: 710–722.

14. Sabeti PC, Varilly P, Fry B, Lohmueller J, Hostetter E, et al. (2007) Genome-wide detection and characterization of positive selection in human populations. Nature 449: 913.

15. Liu N, Zhang K, Zhao H (2008) Haplotype-association analysis. Advances in genetics 60: 335–405.

16. Roach JC, Glusman G, Hubley R, Montsaroff SZ, Holloway AK, et al. (2011) Chromosomal haplotypes by genetic phasing of human families. The American Journal of Human Genetics 89: 382–397.

17. Douglas JA, Boehnke M, Gillanders E, Trent JM, Gruber SB (2001) Experimentally-derived haplotypes substantially increase the efficiency of linkage disequilibrium studies. Nature genetics 28: 361.

18. Ruano G, Kidd KK, Stephens JC (1990) Haplotype of multiple polymorphisms resolved by enzymatic amplification of single DNA molecules. Proceedings of the National Academy of Sciences 87: 6296–6300.

19. Ruano G, Kidd KK (1989) Direct haplotyping of chromosomal segments from multiple heterozygotes via allele-specific PCR amplification. Nucleic acids research 17: 8392.

20. Tininini L, Bertolazzi P, Godi A, Lancia G (2010) CollHaps: a heuristic approach to haplotype inference by parsimony. IEEE/ACM transactions on computational biology and bioinformatics 7: 511–523.

21. Rhee J-K, Li H, Joung J-G, Hwang K-B, Zhang B-T, et al. (2016) Survey of computational haplotype determination methods for single individual. Genes & Genomics 38: 1–12.

22. Wang Y, Feng E, Wang R (2007) A clustering algorithm based on two distance functions for MEC model. Computational biology and chemistry 31: 148–150.

23. Hashemi A, Zhu B, Vikalo H (2018) Sparse tensor decomposition for haplotype assembly of diploids and Polyploids. BMC genomics 19: 191.

24. Olyaee M-H, Khanteymoori A (2018) AROHap: An effective algorithm for single individual haplotype reconstruction based on asexual reproduction optimization. Computational biology and chemistry 72: 1–10.

25. Olyaee M-H, Khanteymoori AR (2019) Single Individual Haplotype Reconstruction Using Fuzzy C-Means Clustering with Minimum Error Correction. Bioinformatics and Biocomputational Research 3.

26. Mazrouee S, Wang W (2014) FastHap: fast and accurate single individual haplotype reconstruction using fuzzy conflict graphs. Bioinformatics 30: i371–i378.

27. Bansal V, Bafna V (2008) HapCUT: an efficient and accurate algorithm for the haplotype assembly problem. Bioinformatics 24: i153–i159.

28. Wang T-C, Taheri J, Zomaya AY (2012) Using genetic algorithm in reconstructing single individual haplotype with minimum error correction. Journal of biomedical informatics 45: 922–930.

29. Patterson M, Marschall T, Pisanti N, Van Iersel L, Stougie L, et al. (2015) WhatsHap: Weighted haplotype assembly for future-generation sequencing reads. Journal of Computational Biology 22: 498–509.

30. Bracciali A, Aldinucci M, Patterson M, Marschall T, Pisanti N, et al. (2016) PWHATSHAP: efficient haplotyping for future generation sequencing. BMC Bioinformatics 17: 342.

31. Bansal V, Halpern AL, Axelrod N, Bafna V (2008) An MCMC algorithm for haplotype assembly from whole-genome sequence data. Genome research 18: 1336–1346.

32. Edge P, Bafna V, Bansal V (2017) HapCUT2: robust and accurate haplotype assembly for diverse sequencing technologies. Genome research 27: 801–812.

33. Chen X, Peng Q, Han L, Zhong T, Xu T (2014) An effective haplotype assembly algorithm based on hypergraph partitioning. Journal of theoretical biology 358: 85–92.

34. Xie M, Wu Q, Wang J, Jiang T (2016) H-PoP and H-PoPG : Heuristic partitioning algorithms for single individual haplotyping of polyploids. Bioinformatics 32: 3735–3744.

35. Puljiz Z, Vikalo H (2016) Decoding genetic variations: Communications-inspired haplotype assembly. IEEE/ACM Transactions on Computational Biology and Bioinformatics (TCBB) 13: 518–530.

36. Cai C, Sanghavi S, Vikalo H (2016) Structured low-rank matrix factorization for haplotype assembly. IEEE Journal of Selected Topics in Signal Processing 10: 647–657.

37. Olyaee MH, Khanteymoori A, Khalifeh K (2019) Application of Chaotic Laws to Improve Haplotype Assembly Using Chaos Game Representation. Scientific reports 9.

38. Mazrouee S, Wang W (2018) PolyCluster: Minimum Fragment Disagreement Clustering for Polyploid Phasing. IEEE/ACM transactions on computational biology and bioinformatics.

39. Han J, Pei J, Yin Y, Mao R (2004) Mining frequent patterns without candidate generation: A frequent-pattern tree approach. Data mining and knowledge discovery 8: 53–87.

40. Han J, Pei J, Yin Y (2000) Mining frequent patterns without candidate generation. ACM sigmod record 29: 1–12.

41. Kuleshov V (2014) Probabilistic single-individual haplotyping. Bioinformatics 30: i379–i385.

42. Barnsley MF (2014) Fractals everywhere: Academic press.

43. Jeffrey HJ (1990) Chaos game representation of gene structure. Nucleic Acids Research 18: 2163–2170.

44. Olyaee MH, Yaghoubi A, Yaghoobi M (2016) Predicting protein structural classes based on complex networks and recurrence analysis. Journal of theoretical biology 404: 375–382.

45. Hoang T, Yin C, Yau SS-T (2016) Numerical encoding of DNA sequences by chaos game representation with application in similarity comparison. Genomics 108: 134–142.

46. Ge L, Liu J, Zhang Y, Dehmer M (2019) Identifying anticancer peptides by using a generalized chaos game representation. Journal of mathematical biology 78: 441–463.

47. Zheng K, Wang L, You Z-H (2019) CGMDA: An Approach to Predict and Validate MicroRNA-Disease Associations by Utilizing Chaos Game Representation and LightGBM. IEEE Access 7: 133314–133323.

48. Anitas EM, Slyamov A (2017) Structural characterization of chaos game fractals using small-angle scattering analysis. PloS one 12.

49. Geraci F (2010) A comparison of several algorithms for the single individual SNP haplotyping reconstruction problem. Bioinformatics 26: 2217–2225.

50. Levy S, Sutton G, Ng PC, Feuk L, Halpern AL, et al. (2007) The diploid genome sequence of an individual human. PLoS biology 5: e254.

51. Chen Z, Fu B, Schweller R, Yang B, Zhao Z, et al. (2008) Linear time probabilistic algorithms for the singular haplotype reconstruction problem from SNP fragments. Journal of Computational Biology 15: 535–546.

52. Zhao Y-Y, Wu L-Y, Zhang J-H, Wang R-S, Zhang X-S (2005) Haplotype assembly from aligned weighted SNP fragments. Computational Biology and Chemistry 29: 281–287.

53. Panconesi A, Sozio M. Fast hare: A fast heuristic for single individual SNP haplotype reconstruction; 2004. Springer. pp. 266–277.

54. Genovese LM, Geraci F, Pellegrini M (2008) SpeedHap: an accurate heuristic for the single individual SNP haplotyping problem with many gaps, high reading error rate and low coverage. IEEE/ACM Transactions on Computational Biology and Bioinformatics (TCBB) 5: 492–502.

55. DePristo MA, Banks E, Poplin R, Garimella KV, Maguire JR, et al. (2011) A framework for variation discovery and genotyping using next-generation DNA sequencing data. Nature genetics 43: 491.

